# Plant Co-expression Annotation Resource: a webserver for identifying targets for genetically modified crop breeding pipelines

**DOI:** 10.1101/2020.05.22.110510

**Authors:** Marcos José Andrade Viana, Adhemar Zerlotini, Mauricio de Alvarenga Mudadu

## Abstract

**Background:** The development of genetically modified crops (GM) includes the discovery of candidate genes through bioinformatics analysis using genomics data, gene expression, and others. Proteins of unknown function (PUFs) are interesting targets for GM crops breeding pipelines for the novelty associated to such targets and also to avoid copyright protections. One method of inferring the putative function of PUFs is by relating them to factors of interest such as abiotic stresses using orthology and co-expression networks, in a guilt-by-association manner.

**Results:** In this regard, we have downloaded, analyzed, and processed genomics data of 53 angiosperms, totaling 1,862,010 genes and 2,332,974 RNA. Diamond and InterproScan were used to discover 72,266 PUFs for all organisms. RNA-seq datasets related to abiotic stresses were downloaded from NCBI/GEO. The RNA-seq data was used as input to the LSTrAP software to construct co-expression networks. LSTrAP also created clusters of transcripts with correlated expression, whose members are more probably related to the molecular mechanisms associated to abiotic stresses in the plants. Orthologous groups were created (OrhtoMCL) using all 2,332,974 proteins in order to associate PUFs to abiotic stress related clusters of co-expression and therefore infer their function in a guilt-by-association manner.

**Conclusion:** A freely available web resource named “Plant Co-expression Annotation Resource” (https://www.machado.cnptia.embrapa.br/plantannot), *Plantannot*, was created to provide indexed queries to search for PUF putatively associated to abiotic stresses. The web interface also allows browsing, querying and retrieving of public genomics data from 53 plants. We hope *Plantannot* to be useful for researchers trying to obtain novel GM crops resistant to climate change hazards.

## BACKGROUND

In the last decades, the ability to genetically engineer plants with success showed the potential to create genetically modified (GM) crops with favourable economic outcomes [1]. As well, in the last decades, the main achievements in this area were genetically improved plants tolerant to herbicide and resistant to insects. Others, like nutritional composition improvements are ongoing [2]. Furthermore, new mechanisms for genome editing are improving the accuracy and speed of genome modifications in plants, such as the CRISPR/CAS system [3,4].

Regarding climate change and environmental factors, plants are being genetically modified to become resilient to abiotic stresses, such as drought, high temperature, rising atmospheric CO2, to potentially overcome the yield losses due to these factors [5,6].

Intellectual property rights (IPR) are vastly used by biotechnology enterprises for their GM plants, to allow exclusive rights and provide better returns for the high investments in research and development [7]. In this way, over the last years many patents applications for genetically improved crops regarding stress tolerance were filled [8].

The first phase for creating GM crops is the candidate gene discovery, which relies on bioinformatics analyses that uses huge volumes of genomics data available on public resources [9,10]. To avoid intellectual property rights over already patented genes, its molecular mechanisms and products, it might be desirable to start researching genes and proteins with no function yet described. These proteins of unknown function (PUF) are very prevalent in eukaryotic genomes and may play roles in determining differences between species [11] and also may be related to resistance to abiotic stresses [12].

Resistance to abiotic stresses is a complex and multigenic trait. Tools and analyses such as QTL, GWAS, gene expression and regulatory networks can be used to find the genes and molecular mechanisms that may play a role in these conditions [13–15] with some results already available [6,16,17].

It is known that differences in the pattern of gene expression, allied to environmental influences, lead to differences in the morphology and phenotype of animals and plants [18]. It is also well established that organs and tissues with the same evolutionary origin have correlated gene expression patterns [19]. To perform molecular comparisons between different species, the focus are genes with the same evolutionary origin, and therefore, with homolog functions, i.e. orthologs [20]. One approach for studying the regulatory functions of a network of genes over different species is to align the co-expression networks using using ortholog genes [21].

In this work we present a web resource named “Plant co-expression annotation resource” (https://www.machado.cnptia.embrapa.br/plantannot) which uses plant genomics and RNA sequencing data, orthology and co-expression networks that allows the selection of PUFs as abiotic stress related candidates to enter GM crop breeding pipelines.

## METHODS

### Raw Data

Genome data (sequence assembly in FASTA formatted files and annotationin GFF files) for 53 angiosperms (Table 1), including *Glycine max* (Gma), *Zea mays* (Zma), *Arabidopsis thaliana* (Ath) and *Oryza sativa* (Osa), were obtained from Phytozome v12 [22] and one from NCBI (*Boea hygrometrica*). The total number of genes and mRNA stored was 1,862,010 and 2,332,974, respectively, together with their translated proteins.

**Table 1:**
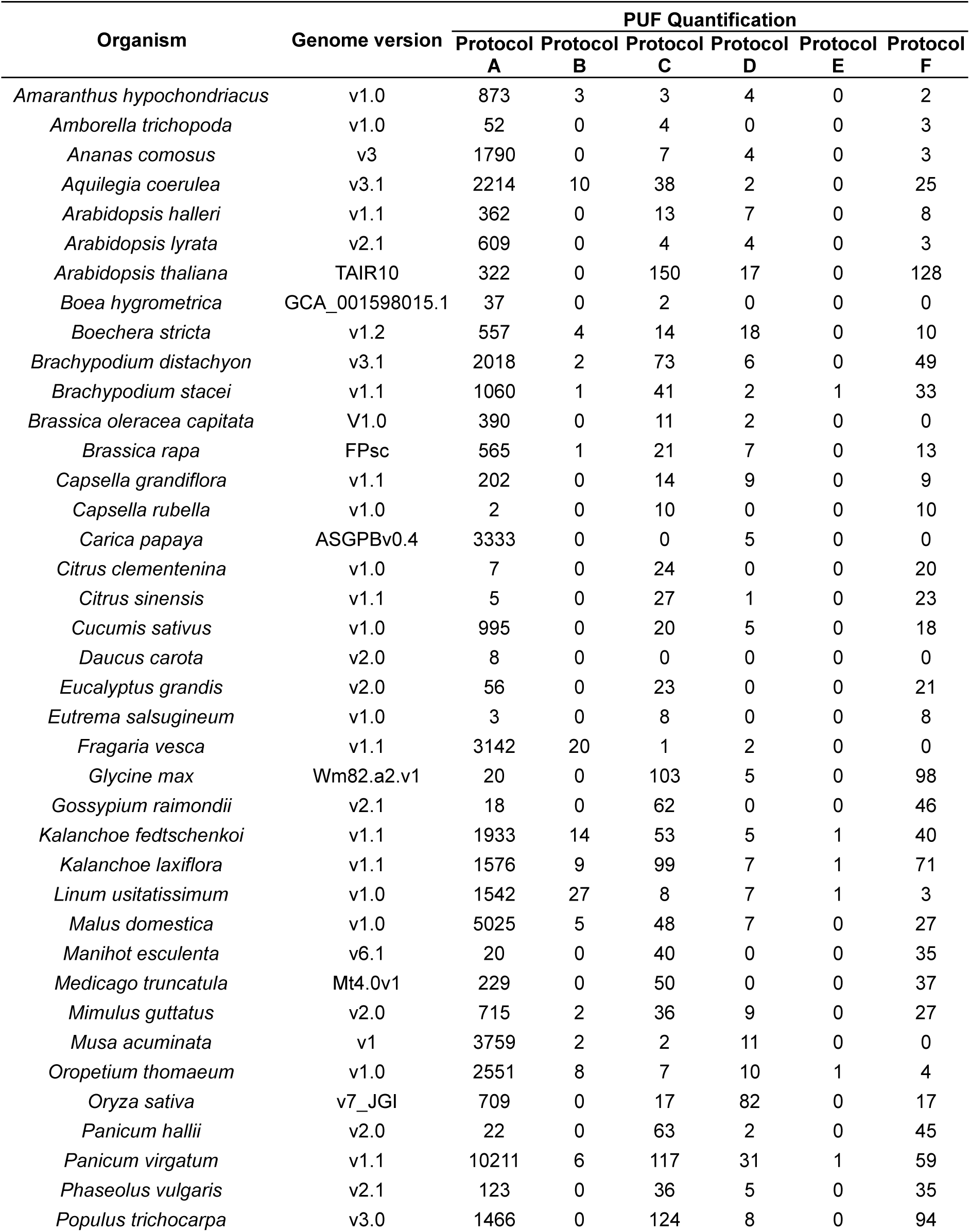
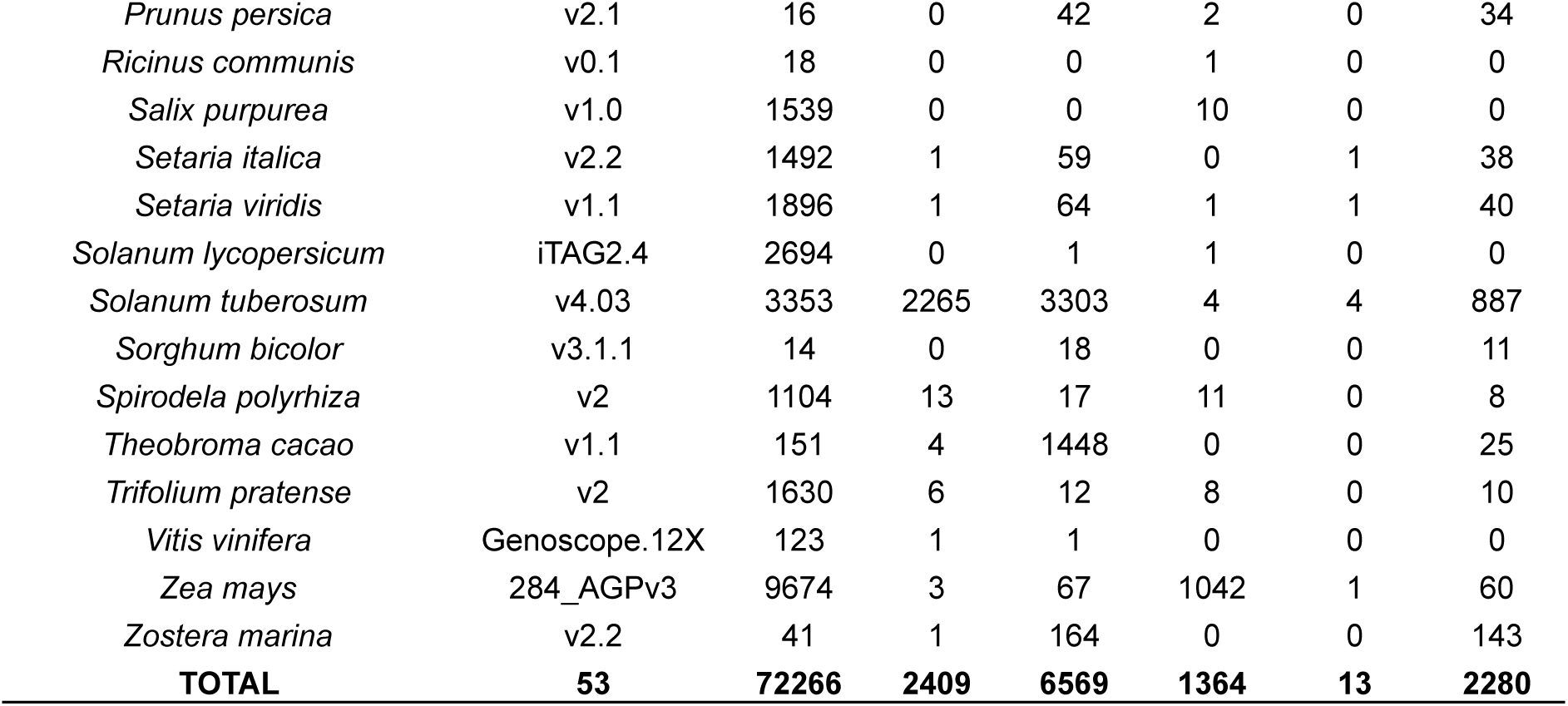
Organisms, genome versions and PUF Quantification.

| Organism | Genome version | PUF Quantification |  |  |  |  |  |
| --- | --- | --- | --- | --- | --- | --- | --- |
|  |  | Protocol A | Protocol B | Protocol C | Protocol D | Protocol E | Protocol F |
| *Amaranthus hypochondriacus* | v1.0 | 873 | 3 | 3 | 4 | 0 | 2 |
| *Amborella trichopoda* | v1.0 | 52 | 0 | 4 | 0 | 0 | 3 |
| *Ananas comosus* | v3 | 1790 | 0 | 7 | 4 | 0 | 3 |
| *Aquilegia coerulea* | v3.1 | 2214 | 10 | 38 | 2 | 0 | 25 |
| *Arabidopsis halleri* | v1.1 | 362 | 0 | 13 | 7 | 0 | 8 |
| *Arabidopsis lyrata* | v2.1 | 609 | 0 | 4 | 4 | 0 | 3 |
| *Arabidopsis thaliana* | TAIR10 | 322 | 0 | 150 | 17 | 0 | 128 |
| *Boea hygrometrica* | GCA_001598015.1 | 37 | 0 | 2 | 0 | 0 | 0 |
| *Boechera stricta* | v1.2 | 557 | 4 | 14 | 18 | 0 | 10 |
| *Brachypodium distachyon* | v3.1 | 2018 | 2 | 73 | 6 | 0 | 49 |
| *Brachypodium stacei* | v1.1 | 1060 | 1 | 41 | 2 | 1 | 33 |
| *Brassica oleracea capitata* | V1.0 | 390 | 0 | 11 | 2 | 0 | 0 |
| *Brassica rapa* | FPsc | 565 | 1 | 21 | 7 | 0 | 13 |
| *Capsella grandiflora* | v1.1 | 202 | 0 | 14 | 9 | 0 | 9 |
| *Capsella rubella* | v1.0 | 2 | 0 | 10 | 0 | 0 | 10 |
| *Carica papaya* | ASGPBv0.4 | 3333 | 0 | 0 | 5 | 0 | 0 |
| *Citrus clementenina* | v1.0 | 7 | 0 | 24 | 0 | 0 | 20 |
| *Citrus sinensis* | v1.1 | 5 | 0 | 27 | 1 | 0 | 23 |
| *Cucumis sativus* | v1.0 | 995 | 0 | 20 | 5 | 0 | 18 |
| *Daucus carota* | v2.0 | 8 | 0 | 0 | 0 | 0 | 0 |
| *Eucalyptus grandis* | v2.0 | 56 | 0 | 23 | 0 | 0 | 21 |
| *Eutrema salsugineum* | v1.0 | 3 | 0 | 8 | 0 | 0 | 8 |
| *Fragaria vesca* | v1.1 | 3142 | 20 | 1 | 2 | 0 | 0 |
| *Glycine max* | Wm82.a2.v1 | 20 | 0 | 103 | 5 | 0 | 98 |
| *Gossypium raimondii* | v2.1 | 18 | 0 | 62 | 0 | 0 | 46 |
| *Kalanchoe fedtschenkoi* | v1.1 | 1933 | 14 | 53 | 5 | 1 | 40 |
| *Kalanchoe laxiflora* | v1.1 | 1576 | 9 | 99 | 7 | 1 | 71 |
| *Linum usitatissimum* | v1.0 | 1542 | 27 | 8 | 7 | 1 | 3 |
| *Malus domestica* | v1.0 | 5025 | 5 | 48 | 7 | 0 | 27 |
| *Manihot esculenta* | v6.1 | 20 | 0 | 40 | 0 | 0 | 35 |
| *Medicago truncatula* | Mt4.0v1 | 229 | 0 | 50 | 0 | 0 | 37 |
| *Mimulus guttatus* | v2.0 | 715 | 2 | 36 | 9 | 0 | 27 |
| *Musa acuminata* | v1 | 3759 | 2 | 2 | 11 | 0 | 0 |
| *Oropetium thomaeum* | v1.0 | 2551 | 8 | 7 | 10 | 1 | 4 |
| *Oryza sativa* | v7_JGI | 709 | 0 | 17 | 82 | 0 | 17 |
| *Panicum hallii* | v2.0 | 22 | 0 | 63 | 2 | 0 | 45 |
| *Panicum virgatum* | v1.1 | 10211 | 6 | 117 | 31 | 1 | 59 |
| *Phaseolus vulgaris* | v2.1 | 123 | 0 | 36 | 5 | 0 | 35 |
| *Populus trichocarpa* | v3.0 | 1466 | 0 | 124 | 8 | 0 | 94 |
| *Prunus persica* | v2.1 | 16 | 0 | 42 | 2 | 0 | 34 |
| *Ricinus communis* | v0.1 | 18 | 0 | 0 | 1 | 0 | 0 |
| *Salix purpurea* | v1.0 | 1539 | 0 | 0 | 10 | 0 | 0 |
| *Setaria italica* | v2.2 | 1492 | 1 | 59 | 0 | 1 | 38 |
| *Setaria viridis* | v1.1 | 1896 | 1 | 64 | 1 | 1 | 40 |
| *Solanum lycopersicum* | iTAG2.4 | 2694 | 0 | 1 | 1 | 0 | 0 |
| *Solanum tuberosum* | v4.03 | 3353 | 2265 | 3303 | 4 | 4 | 887 |
| *Sorghum bicolor* | v3.1.1 | 14 | 0 | 18 | 0 | 0 | 11 |
| *Spirodela polyrhiza* | v2 | 1104 | 13 | 17 | 11 | 0 | 8 |
| *Theobroma cacao* | v1.1 | 151 | 4 | 1448 | 0 | 0 | 25 |
| *Trifolium pratense* | v2 | 1630 | 6 | 12 | 8 | 0 | 10 |
| *Vitis vinifera* | Genoscope.12X | 123 | 1 | 1 | 0 | 0 | 0 |
| *Zea mays* | 284_AGPv3 | 9674 | 3 | 67 | 1042 | 1 | 60 |
| *Zostera marina* | v2.2 | 41 | 1 | 164 | 0 | 0 | 143 |
| **TOTAL** | **53** | **72266** | **2409** | **6569** | **1364** | **13** | **2280** |

RNA-seq data related to abiotic stresses (heat, drought, dehydration and osmotic stress) were downloaded from NCBI/GEO in a total of 17 different GEO Series, 53 GEO Samples and 60 SRA short read files only for Gma, Zma, Gma and Ath (Table 2). The data was obtained by searching GEO datasets for the given organisms using the keywords “stress” and filtering the study type by “Expression profiling by high throughput sequencing”. The raw reads, correspoding to the GEO Samples, were obtained from NCBI/SRA automatically using the sratoolkit v2.9.2 [23].

**Table 2.** GEO experiments, GEO samples and SRA identifiers used to obtain RNA-seq data.

| Organism* | GEO series | GEO samples | SRA | Condition | Tissue | Date |
| --- | --- | --- | --- | --- | --- | --- |
| *Arabidopsis thaliana* | GSE85653 | GSM2280286 | SRR4033018 | Heat stress rep1 | leaves | May-30-2018 |
| *Arabidopsis thaliana* | GSE85653 | GSM2280287 | SRR4033019 | Heat stress rep2 | leaves | May-30-2018 |
| *Arabidopsis thaliana* | GSE85653 | GSM2280288 | SRR4033020 | Heat stress rep3 | leaves | May-30-2018 |
| *Arabidopsis thaliana* | GSE93979 | GSM2466002 | SRR5196729 | WT drought rep1 | leaf | Jun-13-2017 |
| *Arabidopsis thaliana* | GSE93979 | GSM2466003 | SRR5196730 | WT drought rep1 | leaf | Jun-13-2017 |
| *Arabidopsis thaliana* | GSE93420 | GSM2453038 | SRR5167847 | WT_dehydration1 | leaf | Apr-11-2017 |
| *Arabidopsis thaliana* | GSE93420 | GSM2453039 | SRR5167848 | WT_dehydration2 | leaf | Apr-11-2017 |
| *Arabidopsis thaliana* | GSE93420 | GSM2453040 | SRR5167849 | WT_dehydration3 | leaf | Apr-11-2017 |
| *Arabidopsis thaliana* | GSE94015 | GSM2467113 | SRR5197907 | WT RL3h rep1 heat stress (treated at 37C for 3h) | rosette leaves at flower stages 1-9 | Mar-15-2017 |
| *Arabidopsis thaliana* | GSE94015 | GSM2467114 | SRR5197908 | WT RL3h rep2 heat stress (treated at 37C for 3h) | rosette leaves at flower stages 1-9 | Mar-15-2017 |
| *Arabidopsis thaliana* | GSE94015 | GSM2467115 | SRR5197909 | WT RL3h rep3 heat stress (treated at 37C for 3h) | rosette leaves at flower stages 1-9 | Mar-15-2017 |
| *Arabidopsis thaliana* | GSE72806 | GSM1872392 | SRR2302914 | Col h-1R heat stress (44oC for 1h) | leaves | Oct-24-2016 |
| *Arabidopsis thaliana* | GSE72806 | GSM1872393 | SRR2302915 | Col h-2R heat stress (44oC for 1h) | leaves | Oct-24-2016 |
| *Arabidopsis thaliana* | GSE72806 | GSM1872394 | SRR2302916 | Col h-3R heat stress (44oC for 1h) | leaves | Oct-24-2016 |
| *Arabidopsis thaliana* | GSE72806 | GSM1872389 | SRR2302911 | Col s-1R salinity stress | leaves | Oct-24-2016 |
| *Arabidopsis thaliana* | GSE72806 | GSM1872390 | SRR2302912 | Col s-2R salinity stress | leaves | Oct-24-2016 |
| *Arabidopsis thaliana* | GSE72806 | GSM1872391 | SRR2302913 | Col s-3R salinity stress | leaves | Oct-24-2016 |
| *Oryza sativa* | GSE101734 | GSM2714235 | SRR5856930 | Salt | Seedling leaf | Jul-22-2017 |
| *Oryza sativa* | GSE101734 | GSM2714236 | SRR5856931 | Salt | Seedling leaf | Jul-22-2017 |
| *Oryza sativa* | GSE101734 | GSM2714237 | SRR5856932 | Salt | Seedling leaf | Jul-22-2017 |
| *Oryza sativa* | GSE77510 | GSM2053502 | SRR3140959 | Heat stress (45oC) - 12h | leaf | Dec-21-2017 |
| *Oryza sativa* | GSE78972 | GSM2082859 | SRR3209771 | Long Day Drought_S3 | leaf | Mar-01-2017 |
| *Oryza sativa* | GSE78972 | GSM2082860 | SRR3209772 | Long Day Drought_S4 | leaf | Mar-01-2017 |
| *Oryza sativa* | GSE78972 | GSM2082863 | SRR3209775 | Short Day Drought_S7 | leaf | Mar-01-2017 |
| *Oryza sativa* | GSE78972 | GSM2082864 | SRR3209776 | Short Day Drought_S8 | leaf | Mar-01-2017 |
| *Oryza sativa* | GSE78972 | GSM2082866 | SRR3209778 | Long Day Drought_S10 | leaf | Mar-01-2017 |
| *Oryza sativa* | GSE78972 | GSM2082868 | SRR3209780 | Short Day Drought_S12 | leaf | Mar-01-2017 |
| *Oryza sativa* | GSE80811 | GSM2137964 | SRR3466960 | drought - 1 d | leaves | Feb-14-2017 |
| *Oryza sativa* | GSE80811 | GSM2137964 | SRR3466961 | drought - 1 d | leaves | Feb-14-2017 |
| *Oryza sativa* | GSE80811 | GSM2137965 | SRR3466962 | drought - 2 d | leaves | Feb-14-2017 |
| *Oryza sativa* | GSE80811 | GSM2137965 | SRR3466963 | drought - 2 d | leaves | Feb-14-2017 |
| *Oryza sativa* | GSE80811 | GSM2137966 | SRR3466964 | drought - 3 d | leaves | Feb-14-2017 |
| *Oryza sativa* | GSE80811 | GSM2137966 | SRR3466965 | drought - 3 d | leaves | Feb-14-2017 |
| *Oryza sativa* | GSE95668 | GSM2520922 | SRR5311340 | heat - 35oC - 6h | leaf | Nov-07-2017 |
| *Oryza sativa* | GSE95668 | GSM2520923 | SRR5311341 | heat - 35oC - 6h | leaf | Nov-07-2017 |
| *Zea mays* | GSE71723 | GSM1843772 | SRR2144414 | drought | leaf V12 | Feb-04-2016 |
| *Zea mays* | GSE71723 | GSM1843780 | SRR2144422 | drought | leaf V14 | Feb-04-2016 |
| *Zea mays* | GSE71723 | GSM1843788 | SRR2144430 | drought | leaf V16 | Feb-04-2016 |
| *Zea mays* | GSE71723 | GSM1843796 | SRR2144438 | drought | leaf R1 | Feb-04-2016 |
| *Zea mays* | GSE71377 | GSM1833214 | SRR2129983 | drought | leaf | Jan-22-2016 |
| *Zea mays* | GSE71046 | GSM1826061 | SRR2106186 | wt Salt T7 Rep1 | youngest wrapped leaf | Jan-14-2016 |
| *Zea mays* | GSE71046 | GSM1826073 | SRR2106198 | wt Salt T0 Rep2+Rep3 | youngest wrapped leaf | Jan-14-2016 |
| *Zea mays* | GSE71046 | GSM1826077 | SRR2106202 | wt Salt T7 Rep2+Rep3 | youngest wrapped leaf | Jan-14-2016 |
| *Glycine max* | GSE98958 | GSM2628302 | SRR5569810 | dehydrated | leaf | May-31-2018 |
| *Glycine max* | GSE98958 | GSM2628302 | SRR5569811 | dehydrated | leaf | May-31-2018 |
| *Glycine max* | GSE98958 | GSM2628303 | SRR5569812 | dehydrated | leaf | May-31-2018 |
| *Glycine max* | GSE98958 | GSM2628303 | SRR5569813 | dehydrated | leaf | May-31-2018 |
| *Glycine max* | GSE69571 | GSM1704043 | SRR2051086 | salt stress | leaves | Jul-11-2017 |
| *Glycine max* | GSE69571 | GSM1704044 | SRR2051087 | salt stress | leaves | Jul-11-2017 |
| *Glycine max* | GSE69571 | GSM1704045 | SRR2051088 | salt stress | leaves | Jul-11-2017 |
| *Glycine max* | GSE69571 | GSM1704046 | SRR2051089 | salt stress | leaves | Jul-11-2017 |
| *Glycine max* | GSE70310 | GSM1723542 | SRR2079645 | drought (15 days) | leaf r2 stage | Aug-31-2015 |
| *Glycine max* | GSE70310 | GSM1723542 | SRR2079646 | drought (15 days) | leaf r2 stage | Aug-31-2015 |
| *Glycine max* | GSE70310 | GSM1723542 | SRR2079647 | drought (15 days) | leaf r2 stage | Aug-31-2015 |
| *Glycine max* | GSE69469 | GSM1701586 | SRR2048167 | drought (3 days ZT0-8h R1) | leaves v1 stage | Jul-07-2015 |
| *Glycine max* | GSE69469 | GSM1701592 | SRR2048173 | drought (3 days ZT4-12h R1) | leaves v1 stage | Jul-07-2015 |
| *Glycine max* | GSE69469 | GSM1701598 | SRR2048179 | drought (3 days ZT8-16h R1) | leaves v1 stage | Jul-07-2015 |
| *Glycine max* | GSE69469 | GSM1701604 | SRR2048185 | drought (3 days ZT12-20h R1) | leaves v1 stage | Jul-07-2015 |
| *Glycine max* | GSE69469 | GSM1701610 | SRR2048191 | drought (3 days ZT16-24h R1) | leaves v1 stage | Jul-07-2015 |
| *Glycine max* | GSE69469 | GSM1701616 | SRR2048197 | drought (3 days ZT20-4h R1) | leaves v1 stage | Jul-07-2015 |

\* *Gma* (*Glycine max*), *Zma* (*Zea mays*), (*Ath*) *Arabidopsis thaliana* and *Osa* (*Oryza sativa*)

### Analyses

The RNA-seq data was used as input to the *LSTrAP* v1.3 software [15] to construct co-expression networks. Only leaf tissue expression data was used to obtain the networks, to avoid adding noise to the data. LSTrAP was also used to create groups of co-expression, that are clusters of transcripts with correlated expression by using the software MCL version 14-137.

To characterize PUFs, *Diamond v0*.*9*.*24 [24] was used* to align all proteins against the NCBI’s *nr* database (downloaded in January 2018). Diamond BLAST was run with the flag --max-target-seqs 5 and the best hit was selected. *InterproScan* v5.26-65.0 [25] was used to annotate the proteins from the 53 genomes with “Panther” analyses disabled. All other softwares were run using default parameters. Homolog groups were created using *OrhtoMCL* v2.0.9 [26] and the 53 genome’s proteins as input, with default options.

### Framework interface

The *Machado* software [27] was used to store all data and results, and also provide a web server as interface for fast data browsing.

### Filter Protocols

The *Plantannot* software provides several filters and a text search box that allows searching for molecules by its desired annotation features. These filters are needed to obtain PUFs and to try to relate them to abiotic stresses using RNA-seq expression data and co-expression networks. The Filters menu is separated in 8 fields, of those we are going to use only five: “Organism”, “Feature type”, “Orthology”, “Orthologs_coexpression” and “Analyses”. The “Feature Type” filter has three molecule types, from those the polypeptide box is the only that is going to be always checked and the others blank. By using the other 4 remaining filters, 6 protocols were created as examples of different ways to selecting PUFs. Protocol A: using lack of both homology and protein domain signatures. Protocol B: using lack of homology, presence of domain signatures - trying to select Domains of Unknown Function (DUF) from PFAM, and the text search “Unknown function”. Protocol C: using homology, lack of protein domain signatures and the text search “Unknown function”. Protocol D-F: same protocols of A-C but using ortholog groups to find homolog proteins with co-expression data related to abiotic stress. The protocols are explained in Table 3.

**Table 3:** Protocols used to characterize PUFs.

| Name | Objective | Filters (checked boxes only)* |
| --- | --- | --- |
| Protocol A<br>[28] | Find PUFs from organisms whose proteins are not yet in the NCBI's "nr" database and have no protein domain signatures found by InterproScan. | Analyses: no diamond matches.<br>Analyses: no interproscan matches. |
| Protocol B<br>[29] | The same as A but trying to select proteins with the DUF domains from PFAM. | Analyses: no diamond matches.<br>Analyses: interproscan matches.<br>Text search: "Unknown function". |
| Protocol C<br>[30] | Find PUFs from organisms whose proteins are already public in the "nr" database. | Analyses: diamond matches.<br>Analyses: no interproscan matches.<br>Text search: "Unknown function". |
| Protocol D<br>[31] | Same as A but using ortholog groups and co-expression networks to relate proteins to abiotic stress | Analyses: no diamond matches.<br>Analyses: no interproscan matches.<br>Orthology: orthology.<br>Orthologs_coexpression: co-expression. |
| Protocol E<br>[32] | Same as B but using ortholog groups and co-expression networks to relate proteins to abiotic stress | Analyses: no diamond matches.<br>Analyses: interproscan matches.<br>Text search: "Unknown function".<br>Orthology: orthology.<br>ORetryrthologs_coexpression: co-expression. |
| Protocol F<br>[33] | Same as C but using ortholog groups and co-expression networks to relate proteins to abiotic stress | Analyses: diamond matches.<br>Analyses: no interproscan matches.<br>Text search: "Unknown function".<br>Orthology: orthology.<br>Orthologs_coexpression: co-expression. |

\* For all protocols "Feature type: polypeptide" is always checked.

### Availability of Supporting Source Code and Requirements

Project name: Plantannot

Project home page: https://www.machado.cnptia.embrapa.br/plantannot

Operating system(s): Platform independent

Programming language: Python 3; Machado (https://github.com/lmb-embrapa/machado)

Other requirements: None.

License: Free to use

bio.tools id: plantannot

RRID: SCR_018429

## OVERVIEW

An overview of the component processes of the system covering all data and analysis results used as input to the *Machado* framework, can be seen in Figure 1A and are described below in details.

**Figure 1.**
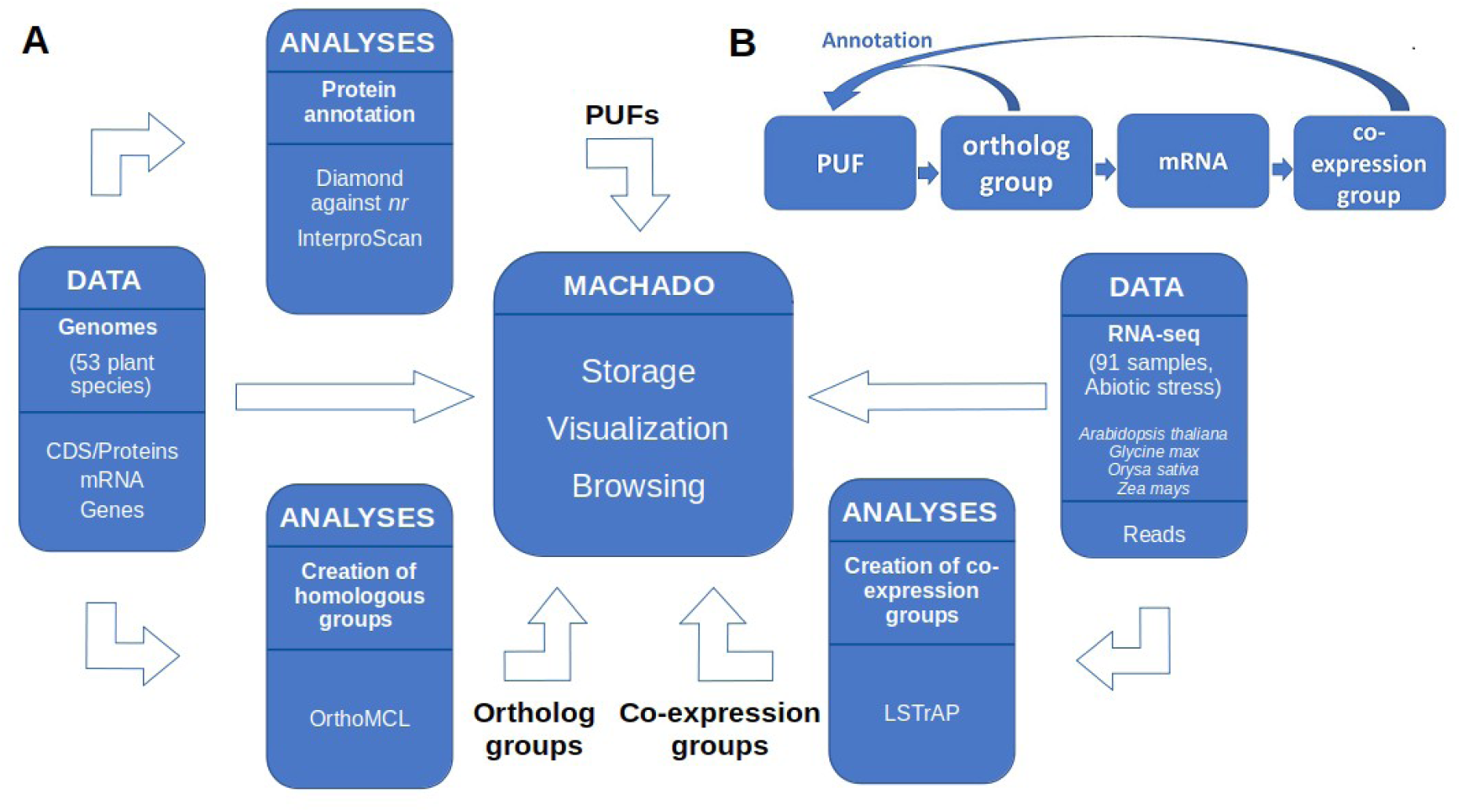
Overview of the Plant Co-expression Annotation Resource processes. B. Guilt-by-association algorithm used to transfer function annotation to PUFs.

### Homolog groups

The 2,332,974 proteins were used as input to the *OrhtoMCL* software to produce 164,267 clusters, or groups of homolog proteins (putative orthologs). All groups comprise 1,900,313 proteins, and the mean cluster size was 11.57 protein members, ranging from 1 to 4,587 members. It is worth mentioning that 8,535 clusters (5,19%) were left with only 1 protein and 75% of all clusters are composed of up to 6 proteins. The ortholog groups are automatically shown in the “Results” frame of the software.

### Co-expression networks

To construct co-expression networks, the 53 GEO Samples (Table 2) were filtered to get expression data only from “leaf” tissue (17, 8, 13 and 15 for Ath, Zma, Gma and Osa respectively). Four co-expression networks were constructed for each of the four organisms (Ath, Zma, Gma and Osa), using the default filters and options of LSTrAP. Groups of co-expression were created using the MCL software following the default instructions in LSTrAP. The MCL software clusters the transcripts with more correlated expression. In this way, the groups of co-expression are supposedly correlated to the molecular mechanisms regarding abiotic stress. 524 groups were obtained (169, 36, 177 and 142 for Ath, Zma, Gma and Osa respectively), with mean size of 140, 113, 282 and 225 for Ath, Zma, Gma e Osa transcript members each, ranging from 1 to 7097 members for Ath, 1 to 4786 for Zma, 1 to 6927 for Gma and 1 to 6636 for Osa.

### PUF characterization

After analyzing all 2,332,974 proteins with *Diamond* and *InterproScan, 72,266* PUFs were characterized (Table 1 – Protocol A) as sequences with no annotation using either *Diamond* or *InterproScan*. Another less sensitive way to find PUFs is to text search for “Unknown proteins” and filter for *InterproScan* matches (e.g.: trying to select PFAM’s DUF domains) only or *Diamond* matches only (e.g.: trying to find proteins with uninformative function annotations), which leads to 2,409 and 6,569 PUFs respectively (Table 1 – Protocols B and C respectively).

### PUF annotation

As there are no information regarding the function of PUFs, one way to infer function is to link PUFs to other molecules by using orthology groups using a guilt-by-association algorithm (Figure 1B). In this way, members from a given ortholog group which already have annotation and/or have protein domains characterized, can be used as a proxy to infer function for the PUF proteins by association. There are 21,895 PUFs as members of ortholog groups which could be a source of functional information and annotation (Protocol A, plus adding the filter “Orthology: orthologs”). Furthermore, whenever a given PUF is part of an ortholog group in which some member, necessarily one of Ath, Gma, Osa or Zma, have its mRNA composing a co-expression group, then by association, the initial PUF is supposedly also related to response to abiotic stresses in plants by inference (see Figure 2). 1364 PUFs were related to co-expression groups using filters that were created to automate this selection (Table 3, Protocol D). This method of searching for PUFs was found to be very strict, since it only retrieves proteins that have no annotations whatsoever. However there are many cases in which PUFs have uninformative annotations, such as: “protein with unknown function”, “putative” or “hypothetical” for example. By modifying Protocol D and text searching for “Unknown function” plus filtering for InterproScan matches only or Diamond matches only, we could annotate 13 and 2,280 PUFs respectively (Table 3, Protocols E and F respectively).

**Figure 2:**
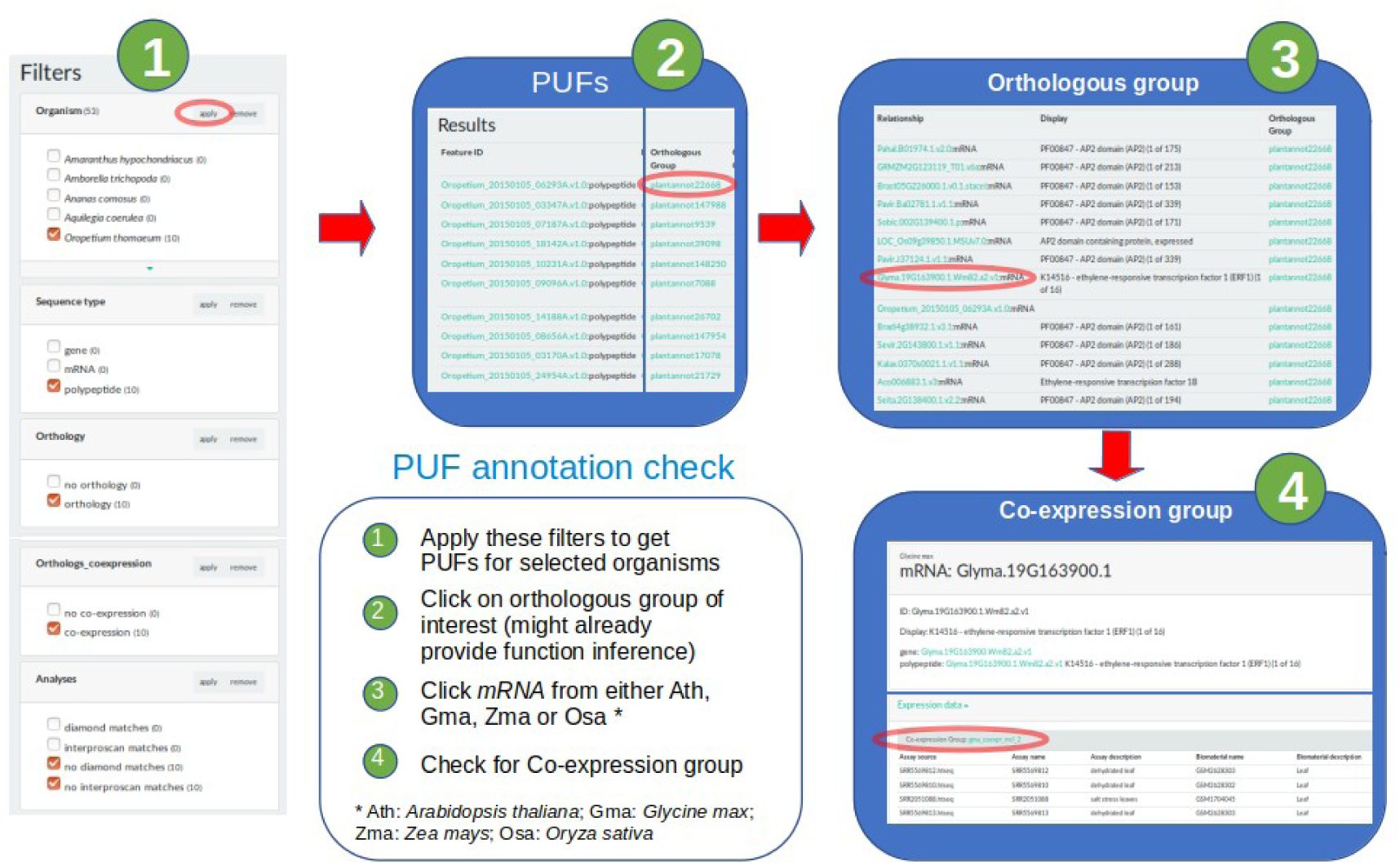
Procotol to check PUF annotation using orthology and co-expression data.

### Case Study: PUF annotations of desiccation tolerant species

We used two species known to be tolerant to desiccation as a pilot study for *Plantannot* as we believe there can be interesting target PUFs related to abiotic stresses to be encountered in these organisms.

#### Oropetium thomaeum

Recently added to the Phytozome database, *Oropetium thomaeum [34]* is a good candidate to discover genes related to abiotic stress. This grass is resilient to extreme and prolonged drying and must have genes involved in the molecular mechanisms related to the control of this phenotype. To find PUFs for *Oropetium thomaeum* one could use Protocol D as described in Table 1. By doing this one will see 10 PUFs in the “Results” page. As there is no annotation for these proteins (although there is one protein that was already annotated as “PTHR13020:SF36 – EXPRESSED PROTEIN (1 of 1” that is not much informative of a function), one can survey the homologous sequences present in the orthologous groups to check for other annotations. In this regard, one can click, for example, on the first member of the “Plantannot22668” group ID, in the “Orthologous Group” column of which the PUF “Oropetium_20150105_06293A.v1.0” is a member. By doing this a new “Results” page will show all members of the “Plantannot22668” group (https://www.machado.cnptia.embrapa.br/plantannot/find/?selected_facets=orthologous_group:plantannot22668). Interestingly the majority of the members are annotated as having an “AP2 domain (PFAM – PF00847)”. By investigating the function of this PFAM domain PF00847 (https://pfam.xfam.org/family/AP2), one can discover that AP2 is a transcription factor that have a major role in hormone regulation [35] and one study shows that there is a binding factor DBF1 that binds AP2 and is related to osmotic stress tolerance and abiotic stress responses in *Arabidopsis thaliana [36]*. By association, it is possible to infer that the PUF “Oropetium_20150105_06293A.v1.0” have a function possibly related to “AP2”, and that orthology could be useful to give novel information for the PUFs. Going further, the “Orthologs_coexpression” box checked before, filtered for orthologous groups of which at least one member participates in a co-expression group. In this way, and also by associative inference, this adds up more evidence that the PUF “Oropetium_20150105_06293A.v1.0” is a good candidate to be related to abiotic stresses and should be further investigated. To check for the co-expression group related to to this PUF, one can follow the procedure in Figure 2 showing that one member of the ortholog group “Plantannot22668” is a protein from Ath, Osa, Zma or Gma, and whose respective mRNA participate in a co-expression group (in this case, the protein from Gma and its mRNA with the same ID: Glyma.19G163900.1.Wm82.a2.v1). This case study can be performed by checking the tutorial session in *Plantannot*’s initial page.

#### Boea hygrometrica (Dorcoceras hygrometricum)

“Drying without dying” is an essential feature in the evolution of earthly plants and *Boea hygrometrica* is an important model of resurrection plant that survives the drying of its leaves and roots without dying [37]. By using a modified version of Protocol F from Table 3 in which we used the text search word “hypothetical”, we recovered 414 PUFs. From these we obtained possible annotations for 199 PUFs (48% of the total) by surveying the orthologous group members as described above. By manually inspecting all 193 annotations we found that 153 (36.95% of the total) had references to abiotic stresses. From these we chose 3 interesting PUFs to describe the possible efficiency of our protocol. The first is the protein KZV45975.1, member of the ortholog group “plantannot11681”, which had members related to “E3 ubiquitin ligase family of proteins”. This family of proteins seems to enhance drought tolerance in *Arabidopsis thaliana* [38]. Another interesting example is the KZV43328.1 protein, member of “plantannot19415” ortholog group, which have 5 members with the PFAM domain “PF00642 - Zinc finger C-x8-C-x5-C-x3-H type (and similar) (zf-CCCH)”. This domain apparently plays roles in abiotic stress response in maize [39]. The final example is the KZV34923.1 protein, who is member of the “plantannot11601” ortholog group which have 17 members that have the PFAM domain “PF05349 - GATA-type transcription activator, N-terminal (GATA-N) (1 of 1)”. It is has been shown that GATA like transcription factors are related to abiotic stress responses in rice [40]. It is worth mentioning that some annotations found refer to abiotic stress that were not part of our RNA-seq data set experimental conditions, like resistance to Aluminum and Cadmium. This could be due to the fact that drought and desiccation tolerance involves a complex process to avoid oxidative damage [41] and we speculate if it may share molecular mechanisms with other kinds of abiotic stresses. The full Boea’s PUF survey can be retrieved from the Supplemental Sheet 1.

## DISCUSSION

Many web servers and online tools available allow navigation and comparative search of expression and co-expression data in plants. Some tools only work online and are not open source like PLAZA 3.0 [42], others are more generic and seek any type of annotation such as CoNeKT [43] and many use also microarray data like the Genevestigator [44]. *Plantannot* has a very specific role of surveying for proteins with unknown function possibly related to abiotic stresses in plants and one of its great differentials is the large number of organisms involved (53 angiosperm species). In addition, the algorithm used to search for PUF annotation includes meta analyses and data relations that involve searches for similarities of sequences, orthology and networks of gene co-expression that are specific and unique.

To demonstrate the potentials of *Plantannot* we devised 6 protocols for filtering sequences of interest. From all the 6 protocols, Protocol A was the most permissive, as it seems that most of the organisms have many proteins that do not return as Diamond best hits against the “nr” database. These sequences were selected by the “no diamond matches” filter and could be retrieved (see table 1). By modifying protocol A and inserting the textual search filter “Unknown function”, led to Protocols B and C.

It is important to mention that genome projects end up having proteins of unknown function annotated in several different ways, by using terms like “hypothetical”, “putative”, “unknown protein”, etc. Therefore, there should be specific text searches for each organism to obtain the best results for selecting PUFs. For example, we needed to adapt the filtering protocols for *Boea hygrometrica*, whose PUFs were best retrieved using the text search “hypothetical”. Other examples can be cited, such as the text search “putative protein” used more efficiently to select PUFs from the organism *Ricinus communis*.

Protocol B uses InterproScan results to search for “Domains of Unknown Function”, or DUFs, from PFAM, that are annotations that could result in more PUFs selected. Protocol C uses the text search to filter Diamond hits and also the original sequence annotations to filter out more PUFs.

The Protocols D-F are more complex protocols that refer to modifications of the Protocols A-C, respectively. They were created by adding filters that could retrieve PUFs that were in the same group of homologous proteins, whose mRNA participate in co-expression network clusters, related do abiotic stresses. This guilt-by-association algorithm explained in Figure 2 led to filtering of many interesting PUFs that would not be highlighted using protocols A-C, such as those described in the study case section.

Protocol D is quite stringent and after applying it, 15 organisms out of 53 involved did not show any results. The reason for this result is that many organisms already have their proteins deposited in the “nr” database and the Diamond best hits would retrieve their own sequence leading them to be filtered out. This occurred with *Boea hygrometrica* but did not occur with *Oropetium tomaeum*, both described in our case studies above.

Many other protocols can still be created, for example, modifying Protocols D-F filtering only by groups of orthologs (filter “Orthology: orthology”) and not by co-expression. This filter selected 21,895 PUFs that belonged to any group of orthologs. This simpler filter could allow one to infer possible functions to these PUFs by just relating them to the annotations found in the members of their common groups of orthologs. Similarly, after applying Protocol D for all organisms, we could manually curate the 1364 PUFs selected, supposedly related to abiotic stress. By conducting a manual search in the groups of orthologs that these PUFs belong, we were able to confirm 159 PUFs with functions possibly related to abiotic stress, found in annotations of ortholog co-members of these PUFs. This result equals 11.6% of the initial PUFs (check the Supplemental Sheet 2 for a complete list of PUFs and annotations for all organisms using this methodology).

## CONCLUSION

We believe that the Plant Co-expression Annotation Resource can be a valuable bioinformatics tool to be used for the search of proof of concept targets to enter pipelines for the creation of genetic modified crops resistant to abiotic stresses and adapted to climate change.

## Supporting information

Supplemental Sheet 1

Supplemental Sheet 2

## AVAILABILITY

The Plant Co-expression Annotation Resource is freely available at https://www.machado.cnptia.embrapa.br/plantannot

## ACKNOWLEDGMENT

Many thanks for Embrapa’s Multiuser Bioinformatics Laboratory (LMB - Laboratório Multiusuário de Bioinformática da Embrapa), UMiP GenClima and Embrapa Agricultural Informatics (Embrapa Informática Agropecuária) for all the support.

